# Serotonergic psychedelic drugs LSD and psilocybin reduce the hierarchical differentiation of unimodal and transmodal cortex

**DOI:** 10.1101/2020.05.01.072314

**Authors:** Manesh Girn, Leor Roseman, Boris Bernhardt, Jonathan Smallwood, Robin Carhart-Harris, R. Nathan Spreng

**Author notes:** To whom correspondence should be addressed. Correspondence, Laboratory of Brain and Cognition, Montreal Neurological Institute and Hospital, 3801 Rue Université, Montréal, QC H3A 2B4. Conflicts of interest: none.

## Abstract

LSD and psilocybin are serotonergic psychedelic compounds with potential in the treatment of mental health disorders. Past neuroimaging investigations have revealed that both compounds can elicit significant changes to whole-brain functional organization and dynamics. A recent proposal linked past findings into a unified model and hypothesized reduced whole-brain hierarchical organization as a key mechanism underlying the psychedelic state, but this has yet to be directly tested. We applied a non-linear dimensionality reduction technique previously used to map hierarchical connectivity gradients to pharmacological resting-state fMRI data to assess cortical organization in the LSD and psilocybin state. Results supported our primary hypothesis: The principal gradient of cortical connectivity, describing a hierarchy from unimodal to transmodal cortex, was significantly flattened under both drugs relative to their respective placebo conditions. Between-condition contrasts revealed that this was driven by a reduction of functional differentiation at both hierarchical extremes – default and frontoparietal networks at the upper end, and somatomotor at the lower. Gradient-based connectivity mapping confirmed that this was underpinned by increased unimodal-transmodal crosstalk. In addition, LSD-dependent principal gradient changes tracked changes in self-reported ego-dissolution. Results involving the second and third gradient, which respectively represent axes of sensory and executive differentiation, also showed significant alterations across both drugs. These findings provide support for a recent mechanistic model of the psychedelic state relevant to therapeutic applications of psychedelics. More fundamentally, we provide the first evidence that macroscale connectivity gradients are sensitive to a pharmacological manipulation, specifically highlighting an important relationship between cortical organization and serotonergic modulation.

## Introduction

The past decade has seen a resurgence of scientific interest in serotonergic psychedelic compounds such as lysergic acid diethylamide (LSD), psilocybin, and dimethyltryptamine (DMT)/ayahuasca, primarily motivated by suggestive findings from preliminary clinical trials for depression, end-of-life-anxiety, alcoholism, and tobacco addiction (Bogenschutz et al., 2015; Carhart-Harris, Bolstridge, et al., 2016; Davis et al., 2020; Gasser, Kirchner, & Passie, 2014; Griffiths et al., 2016; Johnson, Garcia-Romeu, Cosimano, & Griffiths, 2014; Johnson, Hendricks, Barrett, & Griffiths, 2019). Mirroring their complex subjective effects (Preller & Vollenweider, 2016; Schmid et al., 2015; Studerus, Kometer, Hasler, & Vollenweider, 2011), functional neuroimaging investigations have shown significant alterations to whole-brain functional organization and dynamics following psychedelic administration (Carhart-Harris et al., 2012; Carhart-Harris & Friston, 2019; Carhart-Harris, Muthukumaraswamy, et al., 2016; Preller et al., 2018; Preller et al., 2020; Roseman, Leech, Feilding, Nutt, & Carhart-Harris, 2014; Tagliazucchi et al., 2016; Vollenweider & Preller, 2020). More specifically, findings suggest that psychedelic administration shifts the brain towards greater global functional integration, as reflected by the reduced functional segregation of large-scale brain networks and increased global functional connectivity (FC; Carhart-Harris, Muthukumaraswamy, et al., 2016; Müller, Dolder, Schmidt, Liechti, & Borgwardt, 2018; Preller et al., 2018; Roseman et al., 2014; Tagliazucchi et al., 2016). In addition, the complexity of brain dynamics has been shown to increase under psychedelics, as indexed by increases in regional and population-level entropy/complexity (Carhart-Harris, 2018; Lebedev et al., 2016; Schartner, Carhart-Harris, Barrett, Seth, & Muthukumaraswamy, 2017; Varley, Carhart-Harris, Roseman, Menon, & Stamatakis, 2019), as well as increases in the repertoire or dynamic range of functional connectivity (FC) states (Barnett, Muthukumaraswamy, Carhart-Harris, & Seth, 2020; Lord et al., 2019; Luppi et al., 2021; Tagliazucchi, Carhart□Harris, Leech, Nutt, & Chialvo, 2014; Varley et al., 2019)

Notably, the recently proposed RElaxed Beliefs Under Psychedelics (REBUS) model (Carhart-Harris & Friston, 2019) unifies past psychological and neural findings with psychedelics into a common theoretical framework based on hierarchical predictive coding and the Free Energy Principle (Friston, 2010). A primary hypothesis of this model is that psychedelics elicit their characteristic subjective effects by increasing the sensitivity of high-level representations (e.g., beliefs or assumptions) encoded within transmodal cortex to low-level inputs from sensory or limbic sources (Carhart-Harris & Friston, 2019). An acute reduction in the hierarchical differentiation of transmodal versus unimodal cortex, consistent with increased crosstalk between these typically segregated functional zones, would provide support for this hypothesized effect. However, the impact of psychedelics on neural hierarchical organization has yet to be directly tested.

Gradient-mapping techniques have emerged in recent years as valuable tools to characterize brain functional organization (de Wael et al., 2020; Haak, Marquand, & Beckmann, 2018; Huntenburg, Bazin, & Margulies, 2018). In contrast to parcellation approaches which decompose the brain into discrete regions/networks, these approaches model the brain as the superposition of eigenmodes describing continuous axes of differentiation (Haak & Beckmann, 2020; Huntenburg et al., 2018). Such techniques have consistently revealed a principal component describing a gradient of FC spanning from unimodal sensorimotor regions to transmodal association regions centered on the default network (Margulies et al., 2016). This gradient is consistent with tract-tracing work identifying hierarchical cortical organization in human and non-human primates (Mesulam, 1998), corresponds to cortical myeloarchitectonic and transcriptomic transitions (Burt et al., 2018; Huntenburg et al., 2017; Paquola, De Wael, et al., 2019), and represents a functional hierarchy from low-level sensorimotor processing to abstract cognition (Huntenburg et al., 2018; Margulies et al., 2016; Murphy et al., 2018). Gradient-mapping has now been used to characterize neural hierarchy in diverse contexts, including autism (Hong et al., 2019), neonatal development (Larivière et al., 2019), schizophrenia (Dong et al., 2020), and lifespan development (Bethlehem, Paquola, Ronan, et al., 2020; Paquola, Bethlehem, et al., 2019).

The present study investigates alterations of macroscale cortical hierarchy in the psychedelic state by applying gradient-mapping analyses to pharmacological fMRI datasets collected with two serotonergic psychedelic compounds: LSD and psilocybin. We apply cortex-wide and network-specific analyses to characterize psychedelic-dependent changes. In addition, we leverage the organizational scheme provided by the hierarchical gradient to probe hierarchy-specific changes in whole-brain functional connectivity. We predicted that both LSD and psilocybin will be associated with a contraction of the principal gradient, indicating reduced hierarchical differentiation. Further, we predicted that this hierarchical contraction would be specifically consistent with increased crosstalk between unimodal and transmodal cortices. In addition, we investigated psychedelic-induced changes to the second and third gradient, which have been shown, respectively, to differentiate visual from somatomotor/auditory cortex, and executive/task-positive cortex from the rest of cortex. We also predicted gradient dedifferentiation along these axes, given past work which highlighted alterations to sensory and task-positive connectivity in the psychedelic state (Carhart-Harris et al., 2013; Carhart-Harris, Muthukumaraswamy, et al., 2016; Preller et al., 2018; Preller et al., 2020; Roseman et al., 2014). Finally, we examined whether psychedelic-dependent changes in cortical gradients are associated with self-reported subjective effects. Collectively, this study provides the first comprehensive assessment of whether serotonergic drugs LSD and psilocybin pharmacologically alter the primary axes of cortical functional organization. In addition, it constitutes the first direct assessment of whether cortical gradients are sensitive to acute pharmacological manipulation.

## Methods

### Materials and Methods

#### LSD

Neuroimaging data from an already published dataset (Carhart-Harris, Muthukumaraswamy, et al., 2016) was used for the present analyses. The participants and the acquisition protocol are described in greater detail in the original publication and in the Supplementary Information.

In a within-subjects design, resting-state BOLD fMRI data were acquired in 20 subjects for each of LSD and placebo conditions, with conditions spaced 2 weeks apart. The order was counterbalanced across subjects; subjects were blind to this order, but the researchers and those analyzing the data were not. For the placebo condition, subjects were given 10mL of saline and for the LSD condition they were given 75μg of LSD in 10-mL saline via intravenous injection. Three scans were collected on each scan day, 2 without music and 1 without music. The two no-music scans were used in the present analyses (scans 1 and 3). Ego-dissolution scores were collected via intra-scanner visual analogue scale ratings, whereas complex imagery scores were derived from the 11-factor altered states of consciousness (ASC) questionnaire (Dittrich, 1998; Studerus, Gamma, & Vollenweider, 2010) which was completed at the end of each scan day.

#### Psilocybin

Neuroimaging data from an already published dataset (Carhart-Harris et al., 2012) was used for the present analyses. The participants and the acquisition protocol are described in greater detail in the original publication and in the Supplementary Information.

In a within-subjects design, resting-state BOLD fMRI data were acquired in 15 subjects for each of psilocybin and placebo conditions, with conditions spaced 2 weeks apart. The order was counterbalanced across subjects; subjects were blind to this order, but the researchers and those analyzing the data were not. For the placebo condition, subjects were given 10mL of saline and for the psilocybin condition they were given 2mL of psilocybin in 10mL saline, via intravenous injection. Each scanning day consisted of one 12-minute scan with infusion beginning at 6 minutes following the start of the scan. The post-infusion half of the scan for each condition was used in the present analyses. Ego-dissolution scores were collected via intra-scanner visual analogue scale ratings, whereas complex imagery scores were derived from the 11-factor altered states of consciousness (ASC) questionnaire (Dittrich, 1998; Studerus et al., 2010) which was completed at the end of each scan day.

### Ethics Statement

Data collection for both LSD and psilocybin datasets were approved by the National Research Ethics Service committee London-West London and was conducted in accordance with the revised declaration of Helsinki (2000), the International Committee on Harmonization Good Clinical Practice guidelines, and National Health Service Research Governance Framework. Imperial College London sponsored the research, which was conducted under a Home Office license for research with schedule 1 drugs.

### Neuroimaging Data Preprocessing and Denoising

Both datasets underwent an identical preprocessing protocol, as described in detail elsewhere (Carhart-Harris, Muthukumaraswamy, et al., 2016). Subjects with excessive head motion were discarded from analyses (>15% volumes with FD >= 0.5). This resulted in 4 exclusions in the LSD dataset, and 6 exclusions in the psilocybin dataset. The excessive head motion was found in scans conducted during the drug conditions. One additional subject in the LSD dataset exited the scanner due to intra-scanner anxiety, leaving a final sample of 15 for the LSD dataset and 9 for the psilocybin dataset.

The following pre-processing and denoising steps were performed on the BOLD resting-state fMRI data for each dataset: removal of the first three volumes, de-spiking (3dDespike, AFNI), slice time correction (3dTshift, AFNI), motion correction (3dvolreg, AFNI) by registering each volume to the volume most similar to all others, brain extraction (BET, FSL); rigid body registration to anatomical scans, non-linear registration to a 2mm MNI brain (Symmetric Normalization (SyN), ANTS), scrubbing (FD = 0.4), spatial smoothing (FWHM) of 6mm, band-pass filtering between [0.01 to 0.08] Hz, linear and quadratic de-trending (3dDetrend, AFNI), regression of 6 motion parameters, and regression of 3 anatomical nuisance regressors (ventricles, draining veins, and local white matter). Global signal regression was not performed. Quality control tests confirmed the lack of distance-dependent motion confounds in the denoised data (Carhart-Harris et al., 2012; Carhart-Harris, Muthukumaraswamy, et al., 2016).

### Statistical Analysis

#### Gradient-mapping

Cortical gradients were computed using the BrainSpace toolbox (https://github.com/MICA-MNI/BrainSpace; (de Wael et al., 2020)) as implemented in MATLAB. Surfaces were first downsampled from fsaverage 5 space (20,484 vertices) to 10,000 vertices for computational efficiency. For each subject, a 10,000 x 10,000 connectivity matrix was then computed by calculating the pairwise Pearson’s correlation between all vertices. As has been done previously (e.g., Hong et al., 2019; Margulies et al., 2016), this matrix was z-transformed and thresholded row-wise at 90% sparsity in order to retain only the strongest connections. Cosine similarity was then computed on the thresholded z-matrix in order to generate a similarity matrix which captures the similarity in whole-brain connectivity patterns between vertices. This similarity matrix is required as input to the diffusion map embedding algorithm we employed here. The use of cosine similarity as the similarity metric of choice is consistent with past work using this approach (Hong et al., 2019; Margulies et al., 2016; Paquola et al., 2020).

Diffusion map embedding (Coifman et al., 2005; Margulies et al., 2016), a non-linear manifold learning technique from the family of graph Laplacians, was applied to similarity matrices in order to identify gradient components at the individual subject level. The technique estimates a low-dimensional set of embedding components (gradients) from a high-dimensional similarity matrix, where each embedding can intuitively be viewed as a dimension of FC pattern similarity covariance. In the embedding space, vertices that are strongly connected (as weighted by FC pattern similarity) by many connections or a few very strong connections are closer together, whereas vertices with little or no connections are farther apart. Euclidean distance between two points in the embedding space is equivalent to the diffusion distance between probability distributions centered at those points (hence the name of the algorithm), each of which are equivalent to ‘difference in gradient value’ as referred to in the main text. The algorithm is controlled by a single parameter α, which controls the influence of density of sampling points on the manifold (α = 0, maximal influence; α = 1, no influence). Diffusion map embedding is specifically characterized by α = 0.5 (Coifman et al., 2005), which allows the influence of both global and local relationships between data points in the estimation of the embedding space. Following past work (Bethlehem, Paquola, Seidlitz, et al., 2020; Hong et al., 2019), to enable comparisons across subjects, Procrustes rotation was performed to align individual-subject embedding (gradient) components to an all-subjects group average embedding component template. This rotation ensures that gradient axes are matched across subjects. Group contrasts and behavioural associations were conducted using surface-based linear models, as implemented in the SurfStat toolbox (http://www.math.mcgill.ca/keith/surfstat; Worsley et al., 2009).

#### Gradient-based connectivity mapping

In order to further explicate how observed gradient differences related to interregional FC, we examined unimodal versus transmodal functional connectivity changes as stratified by gradient scores. More specifically, we wanted to determine whether unimodal regions (as defined based on the hierarchical gradient) preferentially exhibited increased FC with transmodal regions, as opposed to non-specific global increases. To do this, we first separated vertex-wise gradient scores into 10 percentile bins (0-10, 11-20, 21-30, etc.) for each subject. These 10 bins were then used as ROIs in FC analyses. We were interested in whether each bin preferentially increased its connectivity with unimodal versus transmodal cortex after psychedelic administration. A combined unimodal cortex ROI was created as the combination of the first three percentile bins and a combined transmodal cortex ROI was defined as the last three percentile bins. Unimodal-specific FC and transmodal-specific FC was computed for each of the 10 bins, by computing the Pearson’s correlation (r) between each bin and the combined unimodal ROI and the combined transmodal ROI. T-tests were applied to evaluate drug vs. placebo differences at each bin for each of unimodal-specific and transmodal-specific FC.

To complement this analysis and examine the spatial distribution of FC associated with each gradient percentile bin, we additionally computed seedmaps for each bin and compared across drug and placebo conditions at both vertex-wise and network-wise levels. This enabled a more detailed look at unimodal-specific versus transmodal-specific changes in whole-brain FC.

### Data and code availability

All analyses were conducted in MATLAB using custom code and included functions within the BrainSpace and SurfStat toolboxes described above. Data is freely available at https://openneuro.org/datasets/ds003059/versions/1.0.0.

## Results

We applied gradient-mapping techniques to characterize differences in macroscale cortical gradients in each of LSD and psilocybin states relative to their respective placebo states. The diffusion-map embedding algorithm used here is, compared to other gradient-mapping approaches, robust to noise, computationally inexpensive, and governed by a single parameter controlling the influence of the sampling density on the manifold (see *Methods* for details). Applying this algorithm with standard settings revealed 98 mutually orthogonal gradient components per subject in each of the LSD dataset and psilocybin dataset. We presently include discussion of the first three gradients revealed by this approach, given that these explain the greatest variance and have been highlighted in past work (Margulies et al., 2016).

### Principal gradient

The principal gradient of cortical connectivity revealed in the present datasets replicates past findings (Hong et al., 2019; Margulies et al., 2016) of a putatively hierarchical axis of FC similarity variance spanning from unimodal regions centered in somatomotor cortex on one end to transmodal regions centered on the default network and superior frontal gyrus on the other (Figure 1). This gradient explained the greatest amount of variance in each of LSD placebo (mean 12.9% variance explained), LSD (mean 12.6% variance explained), psilocybin placebo (mean 13.1% variance explained), and psilocybin (mean 11.9% variance explained) conditions.

**Figure 1.**
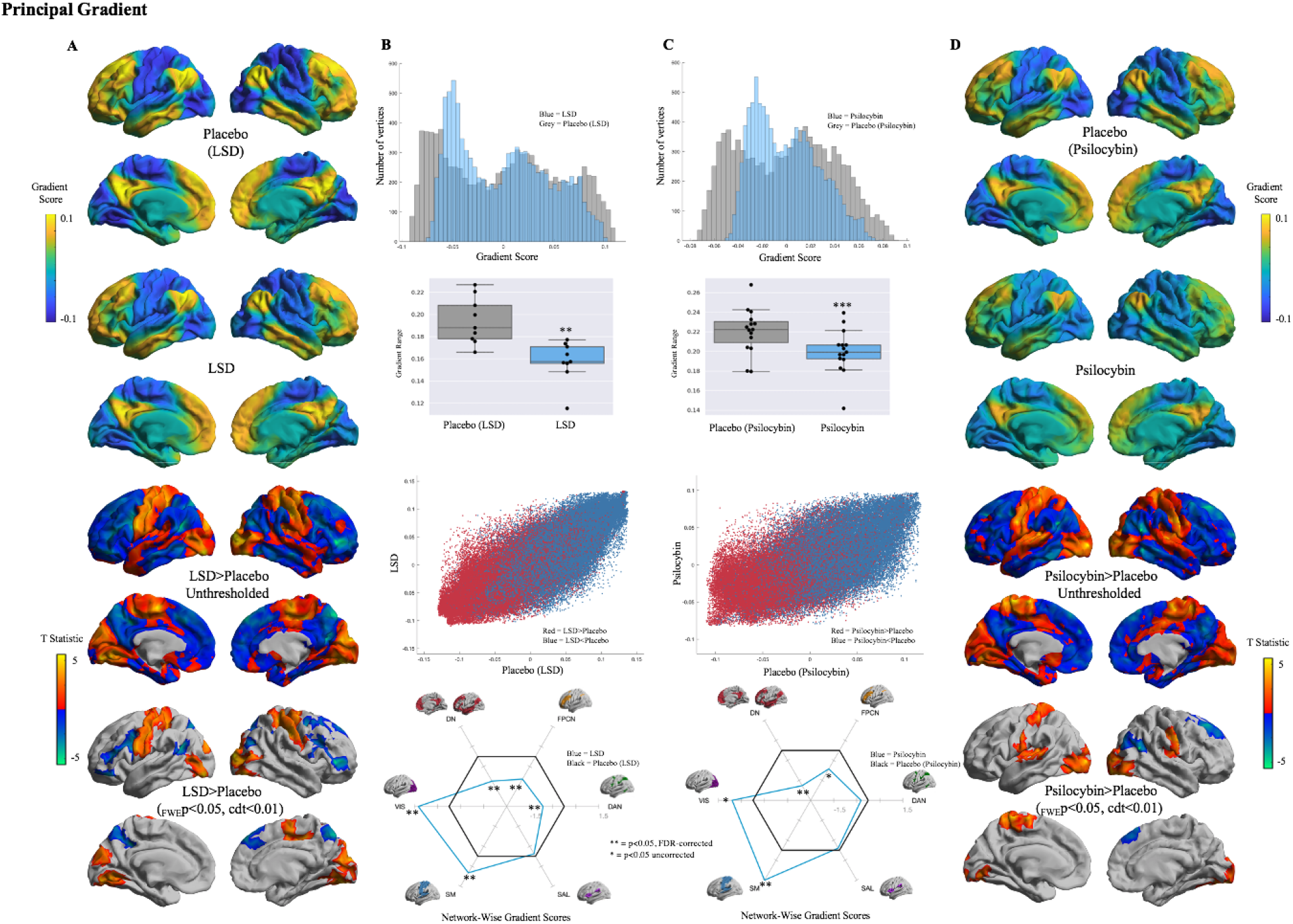
Figures are arranged vertically. **A**. Mean principal gradients representing an axis from unimodal to transmodal cortex, and unthresholded and thresholded vertex-wise contrasts in the LSD dataset **B:** (Top) Histogram showing the distribution of principal gradient values for each of LSD and placebo (LSD) conditions. (Mid-Top) Principal gradient range. Between-condition comparisons indicate a significant contraction in the LSD state (*t*^*28*^ = - 2.5, *p*=0.01). (Mid-Bottom) The principal gradient manifold for both LSD (y-axis) and placebo (LSD) (x-axis) conditions, color coded for overall trends in between-condition differences (see inset). (Bottom) Spider plot displaying mean intra-network principal gradient scores for each of six functional networks, following the (Yeo et al., 2011) parcellation. Values are normalized to the placebo (LSD) condition (black lines). **C:** (Top) Histogram showing the distribution of principal gradient values for each of psilocybin and placebo (psilocybin) conditions. (Mid-Top) Principal gradient range. Between-condition comparisons indicate a significant contraction in the psilocybin state (*t*^*28*^ = -3.9, *p*=0.001). (Mid-Bottom) The principal gradient manifold for both psilocybin (y-axis) an placebo (psilocybin) (x-axis) conditions, color coded for overall trends in between-condition differences (see inset). (Bottom) Spider plot displaying mean intra-network principal gradient scores for each of six functional networks, following the (Yeo et al., 2011) parcellation. Values are normalized to the placebo (psilocybin) condition (black lines). **D:** Mean principal gradients representing an axis from unimodal to transmodal cortex, and unthresholded an thresholded vertex-wise contrasts in the psilocybin dataset. Abbreviations: DN = default network; FPCN = frontoparietal control network; DAN = dorsal attention network; SAL = salience network; SM = somatomotor network; VIS = visual network

Principal gradient histograms for both datasets indicated a contraction on both sides of this gradient in the drug conditions relative to the respective placebo conditions, providing qualitative support for our hypothesis of reduced hierarchical differentiation in the psychedelic state (Figure 1B and 1C Top). To quantitatively assess this, we calculated the difference between each subject’s maximum and minimum principal gradient value and compared these differences across conditions (Figure 1B and 1C Mid-Top). Results confirmed the presence of a significant contraction in both LSD (*t*^*28*^ = -2.5, *p*=0.01) and psilocybin (*t*^*16*^ = -3.9, *p*=0.001) conditions, reflective of reduced functional differentiation along this hierarchical unimodal-transmodal axis. In order to examine overall trends in psychedelic-dependent changes in hierarchical differentiation, we then visualized the principal gradient for each dataset as a scatter plot, color coded for increases (red) and decreases (blue) in each drug condition relative to placebo (Figure 1D). This revealed that both unimodal-proximal regions and transmodal-proximal regions become less differentiated along this axis, suggesting a contraction on both sides of the gradient.

Next, we quantitatively assessed between-condition differences in gradient score values at a vertex-(Figure 1A and 1D Mid-Bottom/Bottom) and network-wise (Figure 1B and 1C Bottom) level. Unthresholded maps at the vertex level are included to display the overall consistency in topology between LSD and psilocybin states, indicating that these distinct serotonergic drugs affect the principal gradient in a similar manner (Figure 1A and 1D Mid-Bottom). LSD vertex-wise contrasts revealed LSD-dependent principal gradient increases in somatomotor cortex, as well as in both medial and lateral visual regions. LSD-dependent principal gradient decreases were found in the precuneus, premotor cortex, superior and inferior frontal gyrus, superior parietal lobule, and Wernicke’s area (_FWE_p<0.05, cdt<0.01). Psilocybin vertex-wise contrasts revealed psilocybin-dependent principal gradient increases in somatomotor and auditory cortex, as well in visual cortex predominantly on the lateral surface. Psilocybin-dependent gradient decreases were found in the superior frontal gyrus and inferior parietal lobule (_FWE_p<0.05, cdt<0.01). Network-wise differences were assessed according to the Yeo et al. (2011) network parcellation scheme. LSD results indicated a significant increase in gradient score within visual and somatomotor networks, and a significant decrease within default, frontoparietal control, and dorsal attention networks (p<0.05 FDR corrected). Psilocybin results indicated a significant increase in gradient score within the somatomotor network and decrease in the default network (p<0.05 FDR corrected). All between-condition differences were consistent with the gradient scores of significant regions and networks approaching zero, indicating reductions in differentiation along this axis. These results therefore offer further quantitative support for the qualitative trend seen in Figure 1B and 1C Mid-Bottom. Namely, that the LSD state is characterized by a pulling-together of both unimodal (visual and somatomotor) and transmodal (default, frontoparietal control, and dorsal attention) networks in gradient space – reflective of relatively symmetrical reduction in differentiation along this hierarchical axis of cortical connectivity. Control analyses indicate that principal gradient score differences are not significantly correlated with motion (Supplementary Figure 1).

### Gradient-Based Connectivity Mapping

In order to further explicate the observed reduction in hierarchical organization in the psychedelic state, we sought to determine whether the observed changes in gradient values were specifically consistent with increased cross-talk between unimodal and transmodal cortices, as hypothesized by the abovementioned REBUS model (Carhart-Harris & Friston, 2019). This interest was further motivated by the fact that the principal gradient changes found in both datasets each represent a movement towards zero. As such, the results are consistent with the changes simply corresponding to less of a loading on this gradient in general and not necessarily to increases/decreases in unimodal- or transmodal-specific connectivity. To evaluate whether the gradient changes are indicative of structured and hierarchically specific changes in FC, we applied a gradient-based connectivity mapping approach. We constructed ROIs based on percentile bins along the principal gradient and evaluated drug-induced unimodal versus transmodal changes in FC. In particular, bin ROIs were correlated with each of a combined unimodal cortex ROI (three lowest bins) and combined transmodal cortex ROI (three highest bins; see Methods for more details).

Between-condition comparisons of unimodal versus transmodal-specific FC supported our hypotheses of increased crosstalk between unimodal and transmodal cortices. For both LSD and placebo, multiple lower percentile bins (corresponding to the unimodal side of the gradient) showed significantly reduced FC with unimodal cortex and less negative FC with transmodal cortex (p<0.05, FDR-corrected or uncorrected; Figure 2). In addition, higher percentile bins (corresponding to the transmodal side of the gradient) displayed significantly increased FC with unimodal cortex but not with transmodal cortex. This suggests that the observed gradient changes have their basis in structured, hierarchically specific changes in FC that are consistent with increased unimodal-transmodal crosstalk.

**Figure 2.**
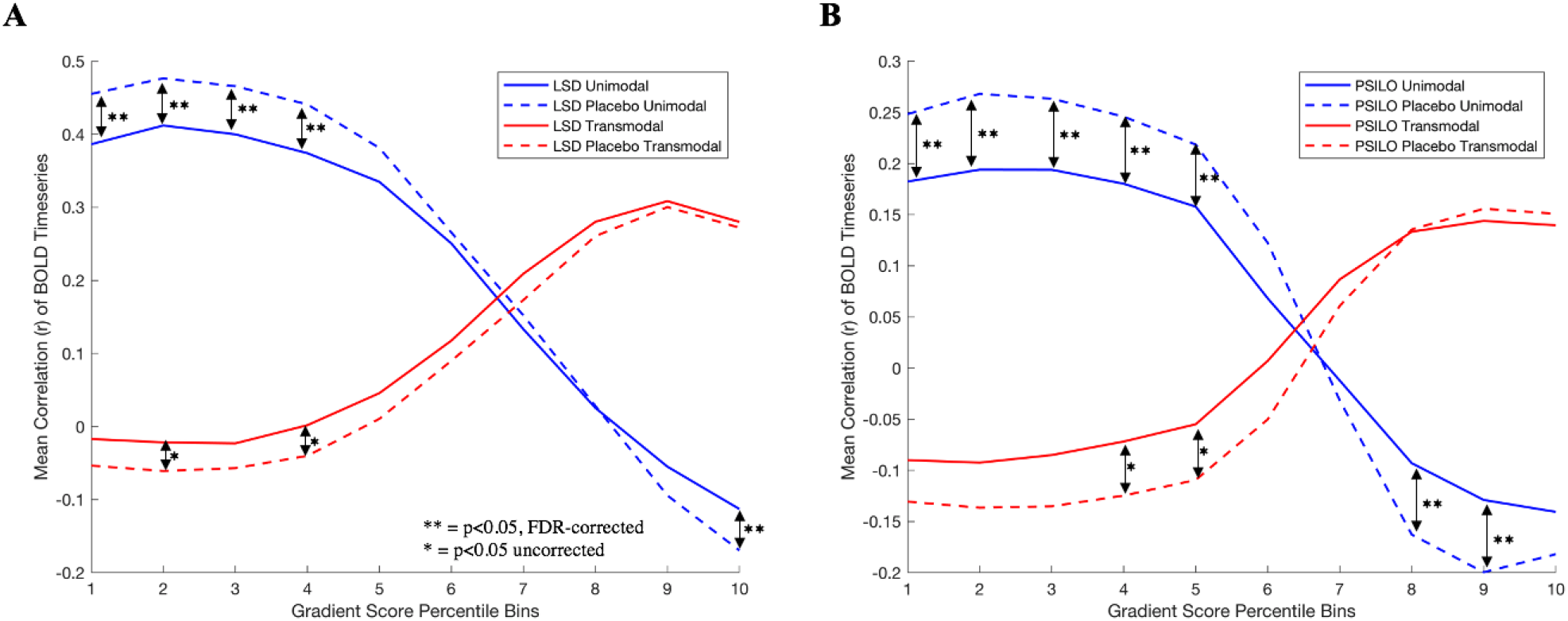
Line plots for each of LSD (A) and psilocybin (B) datasets, indicating unimodal (blue) and transmodal (red) FC for ROIs derived from each of 10 percentile bins along the principal gradient. Solid lines indicate drug conditions, while dotted lines indicate corresponding placebo conditions. ** = p<0.05, FDR-corrected, * = p<0.05, uncorrected.

To further probe this effect, we also examined the spatial distribution of hierarchically specific changes in FC. In particular, we computed whole-brain seedmaps based on each gradient bin ROI and compared across conditions at both the vertex-wise and network-wise level (Figure 3). Results were largely consistent across drugs and indicate that lower percentile bins exhibit significantly reduced FC with somatosensory and visual networks and increased connectivity with the frontoparietal control network, particularly in lateral prefrontal and lateral parietal cortex (_FWE_p<0.05, cdt<0.05 vertex-wise, p<0.05 FDR or uncorrected network-wise). This effect was largely sustained in the median percentile bin, with greater differences involving the default and limbic networks for LSD. With respect to higher percentile bins, with the exception of the default network for the eighth bin of psilocybin, results indicated more of a trend toward increased FC, particularly involving visual, dorsal attention, and salience networks (_FWE_p<0.05, cdt<0.05 vertex-wise, p<0.05 FDR or uncorrected network-wise).

**Figure 3.**
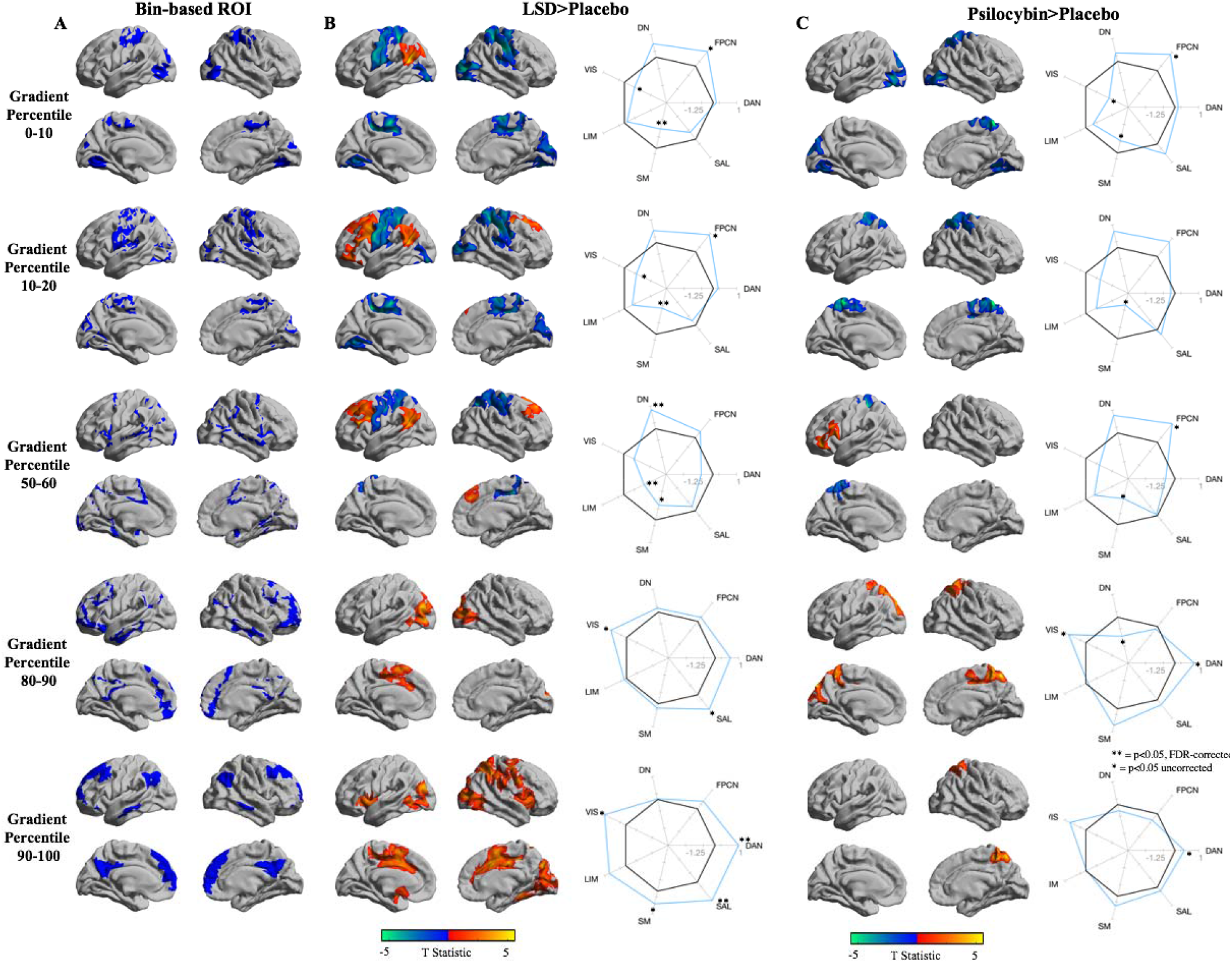
Vertex-wise and network-wise seedmap contrasts for the two lowest, median, and two highest percentile bins. Seeds/ROIs were defined based on percentile bins along the principal gradient. (A) The spatial distribution of each bin-based ROI. (B) LSD>Placebo contrasts: (left) vertex-wise contrast at _FWE_p<0.05, cdt<0.05, (right) network-wise contrast, significance indicated. (C) Psilocybin>Placebo contrasts: (left) vertex-wise contrast at _FWE_p<0.05, cdt<0.05, right network-wise contrast, significance indicated. **=p<0.05, FDR-corrected, *=p<0.05 uncorrected.

### Second and third gradient

Having found support for our hypothesis of a contraction in macroscale functional hierarchy in the LSD state that is consistent with greater unimodal-transmodal crosstalk, we additionally examined the second and third gradient of macroscale functional organization. The second gradient represents an axis of FC similarity variance that separates the visual cortex on one end from the somatosensory/auditory on the other (Figure 4A and 4B). This gradient explained the second most variance in each of LSD placebo (mean 8.8% variance explained), LSD (8.6 variance explained), psilocybin placebo (9.8% variance explained), and psilocybin (mean 10.3% variance explained) conditions. The third gradient represents an axis of FC similarity variance that separates executive control/task-positive regions from the rest of cortex (Figure 4C and 4D). This gradient explained the second most variance in each of LSD placebo (mean 7.7% variance explained), LSD (mean 6% variance explained), psilocybin placebo (mean 8.4% variance explained), and psilocybin (mean 7.5% variance explained) conditions.

**Figure 4.**
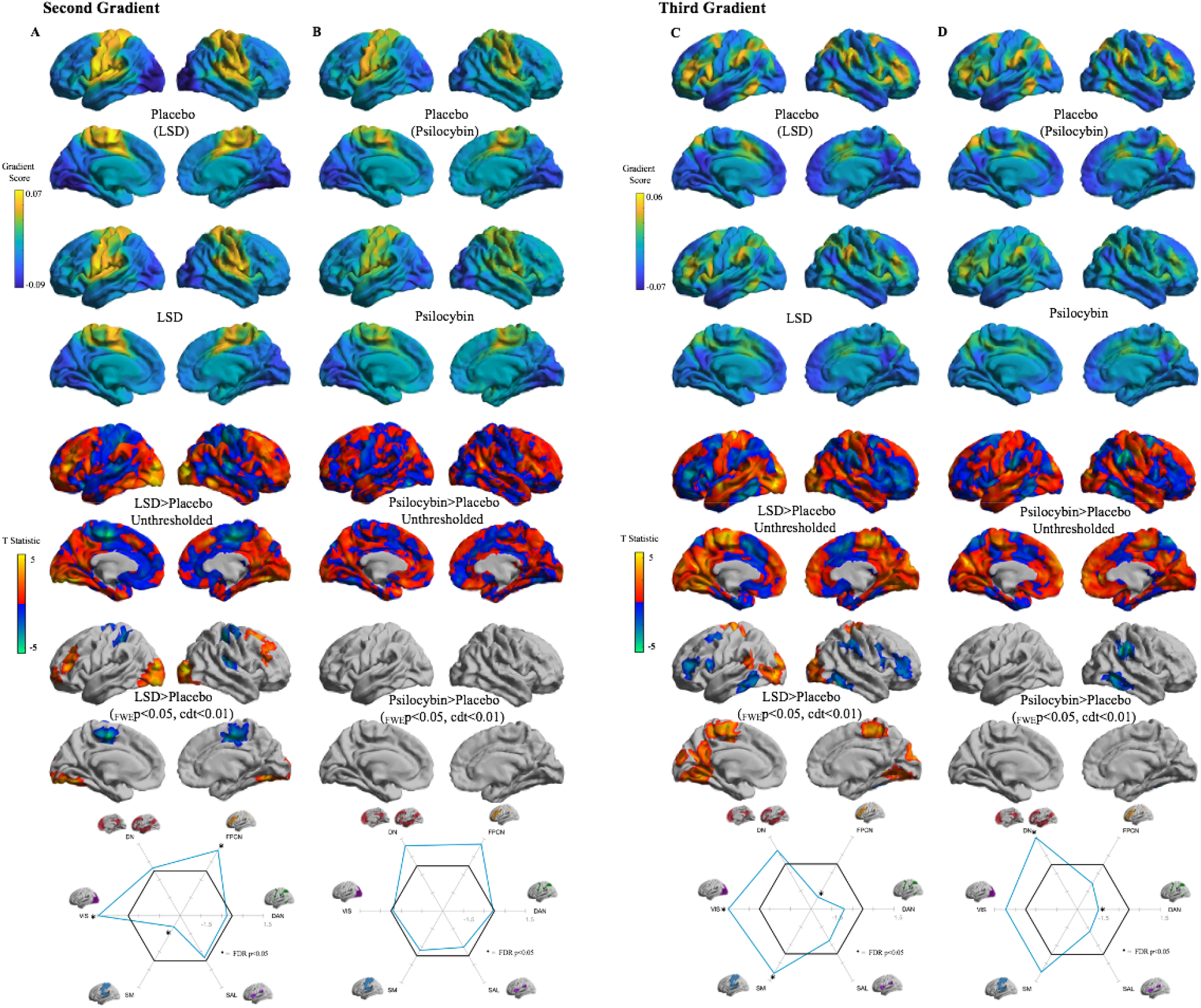
Mean second gradients representing an axis from visual to somatomotor cortex, unthresholded and thresholded vertex-wise contrasts, and network-wise contrasts in the LSD dataset (A) and (B) the psilocybin dataset. (C) Mean third gradients representing an axis from executive control regions to the rest of the cortex, unthresholded and thresholded vertex-wise contrasts, and network-wise contrasts in the LSD dataset and (D) the psilocybin dataset. Network-wise contrasts display mean intra-network gradient values for each of six functional networks, following the (Yeo et al., 2011) parcellation. Values are normalized to the respective placebo conditions (black lines). Blue lines indicate drug condition. Abbreviations: DN = default network; FPCN = frontoparietal control network; DAN = dorsal attention network; SAL = salience network; SMN = somatomotor network; VIS = visual network.

LSD second gradient vertex-wise contrasts revealed LSD-dependent increases in lateral and ventral visual cortex and the middle and superior frontal gyrus bilaterally, while decreases were found in somatomotor and auditory cortex (_FWE_p<0.05, cdt<0.01). Network-wise contrasts revealed significant increases in frontoparietal control and visual networks and decreases in the somatomotor network (FDR p<0.05). Psilocybin vertex- and network-wise contrasts revealed no significant differences, likely due to a lack of power in this dataset. The significant qualitative difference between LSD and the trend-level effects in psilocybin is notable, however, and suggests differential effects on sensory differentiation across these drugs. Visual inspection of the unthresholded contrast maps indicates overlap in topology between psilocybin and LSD-induced changes predominantly within lateral prefrontal cortex and posteromedial cortex. Control analyses indicate that second gradient score differences are not significantly correlated with motion (Supplementary Figure 1).

LSD third gradient vertex-wise contrasts revealed LSD-dependent increases in clusters within somatomotor vortex, lateral and medial visual cortex, and the retrosplenial/posterior cingulate cortex, while decreases were found in bilateral posterior middle temporal gyrus, bilateral inferior frontal gyrus, left premotor cortex, and right supramarginal gyrus (_FWE_p<0.05, cdt<0.01). Network-wise contrasts revealed significant increases in visual and somatomotor networks and decreases in the frontoparietal control network (FDR p<0.05). Psilocybin third gradient vertex-wise contrasts revealed significant decreases in the right supramarginal gyrus and right posterior middle temporal gyrus (_FWE_p<0.05, cdt<0.01). Network-wise contrasts revealed a significant decrease within the dorsal attention network (FDR p<0.05). Control analyses indicate that third gradient score differences are not significantly correlated with motion (Supplementary Figure 1).

### Gradient manifolds

To visualize the relationship between the three gradients examined in this study and how they differ across drug and placebo conditions, we created gradient manifold scatter plots, color coded for each of the Yeo et al. 2011 7 networks (Figure 4). Qualitative examination of the LSD manifold plots (Figure 5A and 5B) reveals an overall less diffuse embedding space distribution in the LSD state, marked by notable contractions on the unimodal aspect of the principal gradient and the visual aspect of the second gradient. Qualitative examination of the psilocybin manifold plots (Figure 5C and 5D) similarly shows a significant contraction on the unimodal side of the principal gradient, but less contraction of the visual network along the second gradient relative to LSD. Psilocybin also displays a greater contraction of transmodal (default and frontoparietal control network) nodes in the principal gradient, and, interestingly, a greater contraction of the third gradient primarily involving regions within salience and dorsal attention networks.

**Figure 5.**
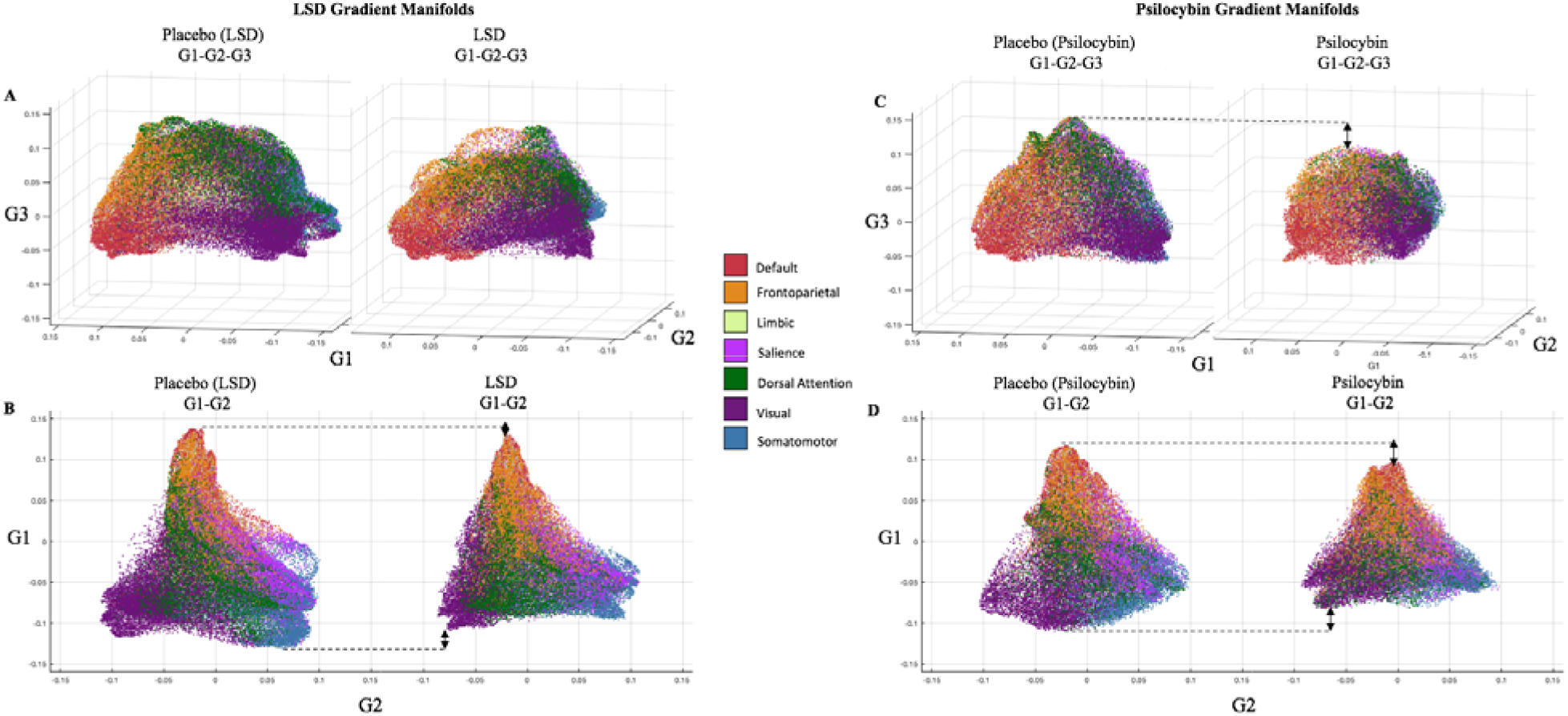
**(A)** Scatterplots representing the gradient embedding space across the first three gradients, for LSD placebo (left) and LSD (right) conditions. **(B)** Scatterplots representing the gradient embedding space across the principal and second gradient, for LSD placebo (left) and LSD (right) conditions. **(C)** Scatterplots representing the gradient embedding space across the first three gradients, for psilocybin placebo (left) and psilocybin (right) conditions. **(D)** Scatterplots representing the gradient embedding space across the principal and second gradient, for psilocybin placebo (left) and psilocybin (right) conditions. Scatter plot colors indicate functional network as per the (Yeo et al., 2011) 7-network parcellation scheme (see inset for legend).. Black arrows indicate contractions along each gradient.

### Relationship to subjective measures

Finally, to ascertain the subjective relevance of changes in macroscale gradients during the psychedelic state, we assessed relationships between drug-placebo gradient differences scores for each drug and each of the three gradients, and drug-placebo differences in two self-report measures which index core components of the psychedelic experience: ego dissolution and complex imagery. Ego dissolution corresponds to the experience that one’s sense of self and what is usually perceived as ‘not self’ (e.g. the physical environment and other individuals) becomes blurred or dissolved completely (Girn & Christoff, 2018; Nour & Carhart-Harris, 2017). Complex imagery corresponds to closed-eye visionary experiences of scenes, landscapes, and images, and is contrasted from simple imagery such as geometric patterns and shapes (Carhart-Harris, Muthukumaraswamy, et al., 2016). One significant relationship was found: a positive association between the principal gradient and ego dissolution in the LSD dataset (_FWE_ *p*<0.05, cdt<0.01; Figure 6). This finding indicates that greater hierarchical differentiation of a large cluster within of the left medial prefrontal cortex/anterior cingulate cortex extending laterally into the superior frontal gyrus was significantly associated with increases in ego dissolution scores. No significant results were found with psilocybin or complex imagery.

**Figure 6.**
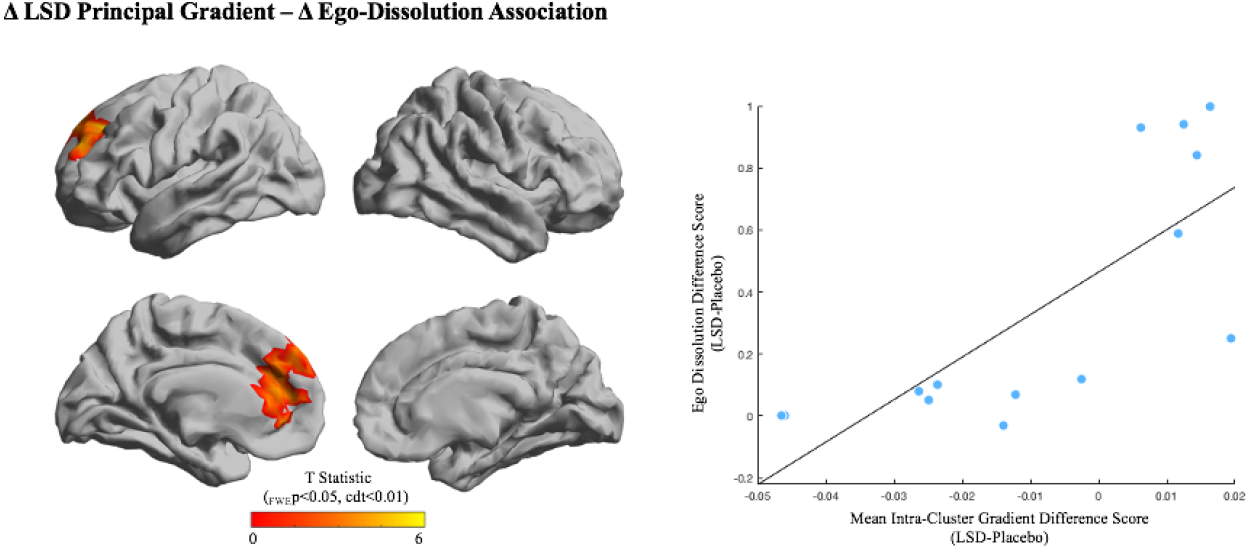
A cluster displaying a significant positive association between LSD-placebo principal gradient difference scores and LSD-placebo self-report difference scores of ego dissolution (_FWE_ p<0.05, cdt<0.01).

## Discussion

To investigate psychedelic-induced changes in cortical functional organization and test the hypothesis that serotonergic psychedelics attenuate brain hierarchical organization, we characterized macroscale cortical gradients after LSD and psilocybin administration. Our results for both LSD and psilocybin datasets replicated past findings (Hong et al., 2019; Margulies et al., 2016) of a principal gradient spanning a hierarchical axis from unimodal to transmodal cortex. Between-condition contrasts supported our primary hypothesis: relative to placebo conditions, this gradient exhibited a significant contraction in both LSD and psilocybin states, reflective of less differentiated hierarchical organization. This was a relatively symmetrical contraction in which somatomotor regions on the unimodal aspect and default and frontoparietal regions on the transmodal aspect both became less differentiated along this hierarchical axis. Gradient-based connectivity mapping indicated that this contraction has its basis in hierarchically specific changes in FC that are consistent with increased crosstalk between unimodal and transmodal cortices. Overall, there was strong convergence in the topographical changes across the LSD and psilocybin datasets, providing important support for the robustness of our findings given th small sample sizes. Results with the second and third gradient indicate alterations to axes of cortical organization related to sensory differentiation and executive region differentiation, respectively. The topography of changes for these two gradients showed strong overlap across datasets, but with greater significant changes with LSD, likely owing to power differences. One divergence was that LSD displayed reduced differentiation of visual and somatomotor cortex along the second gradient, which was not present in psilocybin. These findings extend previous work suggesting greater whole-brain FC of visual cortices in the LSD-state (Carhart-Harris, Muthukumaraswamy, et al., 2016; Preller et al., 2018), a finding that is less consistent with psilocybin (Carhart-Harris, Muthukumaraswamy, et al., 2016; Preller et al., 2020). Finally, we observed significant a significant association between LSD-related changes in the principal gradient and self-reported ego-dissolution – a core component of the psychedelic experience. Collectively, these results provide evidence for significant alterations of macroscale cortical gradients following psychedelic administration, marked by attenuation of hierarchical differentiation between unimodal and transmodal cortices as well as dedifferentiation along sensory and executive axes of cortical organization.

Past neuroimaging investigations with LSD and psilocybin, including past analyses of the present datasets, have revealed that they elicit a complex set of changes to both static and dynamic FC, as well as to the entropy/complexity of regional timeseries (Barnett et al., 2020; Carhart-Harris, 2018; Carhart-Harris, Muthukumaraswamy, et al., 2016; Lebedev et al., 2016; Lord et al., 2019; Luppi et al., 2021; Müller et al., 2018; Preller et al., 2018; Roseman et al., 2014; Schartner et al., 2017; Tagliazucchi et al., 2014; Tagliazucchi et al., 2016; Varley et al., 2019). This work has often attempted to manage this complexity by describing results either in terms of a focus on specific large-scale network interactions (e.g., involving the default network) or in terms of general trends (e.g., towards global integration; Carhart-Harris, Muthukumaraswamy, et al., 2016; Preller et al., 2018; Preller et al., 2020; Roseman et al., 2014; Tagliazucchi et al., 2016). This approach, while useful and necessary given the infancy of the field, can obscure important nuances in the structure of the data. For example, although decreased within-network and increased between-network FC is often listed as a consistent finding with serotonergic psychedelics, the effects with respect to specific networks/network pairs has limited overlap across drugs and datasets (Carhart-Harris, Muthukumaraswamy, et al., 2016; Müller et al., 2018; Roseman et al., 2014). Although this is to be somewhat expected given small sample sizes and analytical discrepancies in this area of research, it suggests the need for approaches which can help organize findings and facilitate comparisons. The cortical gradient approach used here provides a novel perspective on the neural effects of serotonergic psychedelics by collapsing the complex mosaic of increases and decreases in FC into a low-dimensional set of macroscale axes which suggest specific alterations in cortical information processing. This approach can facilitate both functional interpretation and the identification of effects specific to this class of drugs, which, evidently, decreased network segregation is not. As an example, healthy aging is also characterized by decreased within-network and increased between-network FC, yet gradient analyses comparing young and older adults have revealed qualitatively different effects than those observed here (Bethlehem, Paquola, Seidlitz, et al., 2020; Setton et al., 2021).

The greatest LSD-dependent increases in the unimodal-transmodal hierarchical gradient were in regions at the lowest end of the cortical hierarchy – i.e., the somatomotor cortex, while the greatest decreases occurred within regions at the highest end – i.e., regions pertaining to the default and frontoparietal control networks such as the ventral posterior cingulate/retrosplenial cortex, the precuneus, and the superior frontal gyrus. This finding provides evidence for a relatively symmetrical contraction of hierarchical organization under LSD. Our gradient-based connectivity mapping approach further indicated that this contraction was underpinned by hierarchically specific changes in FC. In particular, our results revealed that regions on the unimodal side of the brain’s hierarchy become less connected amongst themselves and more connected to transmodal regions, while regions on the transmodal side of the hierarchy maintain their connections amongst themselves and become more connected to unimodal regions. This suggests the movement of the brain into a more integrative mode of functioning, wherein unimodal regions are less modular and less differentiated from the globally distributed network of transmodal hubs. Moreover, the reduction in hierarchical differentiation and increased unimodal-transmodal crosstalk observed here is consistent with a reduction in the number of intervening processing steps between low-level sensorimotor cortex and high-level association cortex. This can be understood in the context of resting-state FC investigations of cortical organization which have provided evidence that cortical signals propagate through the macroscale processing hierarchy moving from modular sensory processing, to multi-modal integration, to higher-order processing within a distributed network of transmodal hubs (Hong et al., 2019; Paquola et al., 2020; Park et al., 2020; Sepulcre, Sabuncu, Yeo, Liu, & Johnson, 2012; Vazquez-Rodriguez, Liu, Hagmann, & Misic, 2020). In addition, both meta-analytic and task-based analyses have provided evidence that lower levels of the hierarchy pertain to behaviours that are coupled to immediate sensory input, while higher hierarchical zones pertain to perceptually-decoupled, abstract cognitive processes (Margulies et al., 2016; Mckeown et al., 2020; Murphy et al., 2018). Thus, in the psychedelic state, the observed levelling of hierarchy may correspond to decreased functional differentiation between sensory and abstract cognitive processing – consistent with the predictions of the REBUS model (Carhart-Harris & Friston, 2019).

This notion of dedifferentiation between concrete and abstract processing is consistent with past reports suggesting that psychedelics can elicit a blurring of the internal-external/subject-object distinction alongside an increased influence of internal mentation on perceptual processing (K. C. R. Fox, Girn, Parro, & Christoff, 2018; Girn & Christoff, 2018; Girn, Mills, Roseman, Carhart-Harris, & Christoff, 2020; Kraehenmann et al., 2017; Millière, 2017). Of particular interest is the phenomenon of ‘ego-dissolution’, which is a central component of the psychedelic experience and is associated with so-called ‘mystical-type experiences’ which are core predictors of the therapeutic efficacy of psychedelic treatment approaches (Johnson et al., 2019). It corresponds to an experience of blurred or dissolved boundaries between self and world/other and a reduction in the perceived centrality of one’s personal problems and concerns (Girn & Christoff, 2018; Millière, 2017; Nour, Evans, Nutt, & Carhart-Harris, 2016). We observed a significant positive association between LSD principal gradient scores and ego-dissolution within left medial prefrontal cortex/anterior cingulate cortex (mPFC/ACC) – a region that has been consistently implicated in self-related processing (D’Argembeau et al., 2007; Dixon, Thiruchselvam, Todd, & Christoff, 2017). This suggests a potential relationship between mPFC/ACC hierarchical differentiation and ego-dissolution. It is important to note, however, that replication in a larger sample is needed to ascertain the reliability of this effect given the small sample size and lack of replication in the psilocybin dataset. Nonetheless, this finding suggests the potential utility of characterizing changes in unimodal-transmodal differentiation for revealing novel associations between brain function and clinically-relevant dimensions of the psychedelic experience.

In addition to changes in cortical hierarchy, we examined changes in the second and third gradient of cortical connectivity. The second gradient represents the differentiation of visual versus somatomotor/auditory processing. We found multiple changes in this gradient with LSD which were not present with psilocybin. Most notable, there was a significant reduction in visual network differentiation along this axis which was trending in the opposite direction with psilocybin. This is consistent with past work that has shown significant increases in V1 whole-brain FC in the LSD state (Carhart-Harris, Muthukumaraswamy, et al., 2016; Preller et al., 2018), whereas similar changes are less consistent with psilocybin (Carhart-Harris et al., 2012; Preller et al., 2020). The third gradient represents an axis which differentiates the so-called task positive (M. D. Fox et al., 2005)/multiple demand (Duncan, 2010)/executive network (Niendam et al., 2012; Seeley et al., 2007) from the rest of the cortex. Results with both drugs broadly indicate a contraction of this gradient consistent with reduced differentiation of executive regions from the rest of cortex. Although speculative, this might relate to certain cognitive impairments present while under the influence of these drugs (Pokorny, Duerler, Seifritz, Vollenweider, & Preller, 2020).

Finally, to our knowledge, this is the first investigation which has shown significant pharmacologically-induced alterations of macroscale cortical gradients. In particular, we have demonstrated that cortical gradients can be acutely altered by two distinct serotonergic manipulations and, further, that these changes may be paralleled by specific changes in conscious experience. Although LSD and psilocybin exhibit a complex pharmacology, past work has robustly linked their characteristic ‘psychedelic’ effects and alterations to whole-brain functional dynamics to partial agonism at the 5-HT2A receptor (Kraehenmann et al., 2017; Nichols, 2016; Preller et al., 2018; Preller et al., 2017). These findings therefore shed novel light on a potential role for the 5-HT2A receptor in modulating the functional differentiation of unimodal and transmodal systems, as well as in modulating macroscale axes of cortical connectivity more generally. More research is needed to investigate serotonergic influence on global brain dynamics.

An important limitation to the present results is the relatively small sample sizes. For this reason, we encourage cautious interpretation of our findings. However, the convergence across drugs found here provides strong evidence for the robustness of the observed effects. Despite consistent evidence suggesting their safety in controlled research settings (Johnson, Richards, & Griffiths, 2008; Schmid et al., 2015), collecting large datasets with psychedelics is currently difficult due to hurdles pertaining to funding and ethics board approval (Nutt, King, & Nichols, 2013). We hope our findings serve as a foundational guide for future research in this nascent field and as motivation for replication in larger samples.

The present study extends past findings on the neural underpinning of the psychedelic state by revealing that the whole-brain effects of LSD and psilocybin can be represented as a contraction the brain’s macroscale functional hierarchy – directly in line with a recently proposed unified model of psychedelic brain action (Carhart-Harris & Friston, 2019). We also provided evidence of reductions in sensory and executive region differentiation following psychedelic administration. Future work is needed to ascertain whether reductions in cortical hierarchy in the acute psychedelic experience directly relate to findings of rapid and sustained symptom reductions observed via psychotherapeutically-mediated experiences with these drugs (Johnson et al., 2019). The findings of this study lend further weight to the view that psychedelics can be powerful tools for investigating brain organization and dynamics.

## Acknowledgements

MG acknowledges funding from National Sciences and Engineering Research Council of Canada (NSERC; Alexander Graham Bell Canada Graduate Scholarships, CGS-D). JS is supported by a European Research Council Consolidator award (WANDERINGMINDS– 646927). BB acknowledges research funding from the Canadian Institutes of Health Research (CIHR FDN-154298), the National Sciences and Engineering Research Council of Canada (NSERC; Discovery-1304413), and the Canada Research Chairsprogram (CRC-Tier 2 in Cognitive Neuroinformatics of Healthy and Diseased Brains). RLC-H is supported by the Alex Mosley Charitable Trust and supporters of the Centre for Psychedelic Research: https://www.imperial.ac.uk/psychedelic-research-centre. The original study received support from a Crowd Funding Campaign and the Beckley Foundation, as part of the Beckley-Imperial Research Programme.

## References

Barnett, L., Muthukumaraswamy, S. D., Carhart-Harris, R. L., & Seth, A. K. (2020). Decreased directed functional connectivity in the psychedelic state. Neuroimage, 209, 116462.

Bethlehem, R. A., Paquola, C., Ronan, L., Seidlitz, J., Bernhardt, B., & Tsvetanov, K. A. (2020). Dispersion of functional gradients across the lifespan. bioRxiv.

Bethlehem, R. A., Paquola, C., Seidlitz, J., Ronan, L., Bernhardt, B., Tsvetanov, K. A., & Consortium, C.-C. (2020). Dispersion of functional gradients across the adult lifespan. Neuroimage, 222, 117299.

Bogenschutz, M. P., Forcehimes, A. A., Pommy, J. A., Wilcox, C. E., Barbosa, P., & Strassman, R. J. (2015). Psilocybin-assisted treatment for alcohol dependence: A proof-of-concept study. Journal of Psychopharmacology, 29(3), 289–299.

Burt, J. B., Demirtaş, M., Eckner, W. J., Navejar, N. M., Ji, J. L., Martin, W. J., … Murray, J. D. (2018).Hierarchy of transcriptomic specialization across human cortex captured by structural neuroimaging topography. Nature Neuroscience, 21(9), 1251–1259.

Carhart-Harris, R. L. (2018). The entropic brain-revisited. Neuropharmacology, 142, 167–178.

Carhart-Harris, R. L., Bolstridge, M., Rucker, J., Day, C. M., Erritzoe, D., Kaelen, M., … Feilding, A. (2016). Psilocybin with psychological support for treatment-resistant depression: an open-label feasibility study. The Lancet Psychiatry, 3(7), 619–627.

Carhart-Harris, R. L., Erritzoe, D., Williams, T., Stone, J. M., Reed, L. J., Colasanti, A., … Murphy, K. (2012). Neural correlates of the psychedelic state as determined by fMRI studies with psilocybin. Proceedings of the National Academy of Sciences, 109(6), 2138–2143.

Carhart-Harris, R. L., & Friston, K. J. (2019). REBUS and the Anarchic Brain: Toward a Unified Model of the Brain Action of Psychedelics. Pharmacological Reviews, 71(3), 316–344.

Carhart-Harris, R. L., Leech, R., Erritzoe, D., Williams, T. M., Stone, J. M., Evans, J., … Nutt, D. J. (2013). Functional connectivity measures after psilocybin inform a novel hypothesis of early psychosis. Schizophrenia bulletin, 39(6), 1343–1351.

Carhart-Harris, R. L., Muthukumaraswamy, S., Roseman, L., Kaelen, M., Droog, W., Murphy, K., … Nutt, D. J. (2016). Neural correlates of the LSD experience revealed by multimodal neuroimaging. Proceedings of the National Academy of Sciences, 113(17), 4853-4858. doi:10.1073/pnas.1518377113

Coifman, R. R., Lafon, S., Lee, A. B., Maggioni, M., Nadler, B., Warner, F., & Zucker, S. W. (2005). Geometric diffusions as a tool for harmonic analysis and structure definition of data: Diffusion maps. Proceedings of the National Academy of Sciences, 102(21), 7426–7431.

D’Argembeau, A., Ruby, P., Collette, F., Degueldre, C., Balteau, E., Luxen, A., … Salmon, E. (2007).Distinct regions of the medial prefrontal cortex are associated with self-referential processing and perspective taking. Journal of Cognitive Neuroscience, 19(6), 935–944.

Davis, A. K., Barrett, F. S., May, D. G., Cosimano, M. P., Sepeda, N. D., Johnson, M. W., … Griffiths R. R. (2020). Effects of psilocybin-assisted therapy on major depressive disorder: a randomized clinical trial. JAMA Psychiatry.

de Wael, R. V., Benkarim, O., Paquola, C., Lariviere, S., Royer, J., Tavakol, S., … Valk, S. (2020).BrainSpace: a toolbox for the analysis of macroscale gradients in neuroimaging and connectomics datasets. Communications biology, 3(1), 1–10.

Dittrich, A. (1998). The standardized psychometric assessment of altered states of consciousness (ASCs) in humans. Pharmacopsychiatry.

Dixon, M. L., Thiruchselvam, R., Todd, R., & Christoff, K. (2017). Emotion and the prefrontal cortex: An integrative review. Psychological Bulletin, 143(10), 1033.

Dong, D., Luo, C., Guell, X., Wang, Y., He, H., Duan, M., … Yao, D. (2020). Compression of Cerebellar Functional Gradients in Schizophrenia. Schizophrenia bulletin.

Duncan, J. (2010). The multiple-demand (MD) system of the primate brain: mental programs for intelligent behaviour. Trends in Cognitive Sciences, 14(4), 172–179.

Fox, K. C. R., Girn, M., Parro, C., & Christoff, K. (2018). Functional neuroimaging of psychedelic experience: An overview of psychological and neural effects and their relevance to research on creativity, daydreaming, and dreaming. The cambridge handbook of the neuroscience of creativity, 92–113.

Fox, M. D., Snyder, A. Z., Vincent, J. L., Corbetta, M., Van Essen, D. C., & Raichle, M. E. (2005). The human brain is intrinsically organized into dynamic, anticorrelated functional networks. Proc Natl Acad Sci U S A, 102(27), 9673–9678.

Friston, K. J. (2010). The free-energy principle: a unified brain theory? Nature Reviews Neuroscience, 11(2), 127.

Gasser, P., Kirchner, K., & Passie, T. (2014). LSD-assisted psychotherapy for anxiety associated with a life-threatening disease: A qualitative study of acute and sustained subjective effects. Journal of Psychopharmacology, 29(1), 57–68.

Girn, M., & Christoff, K. (2018). Expanding the Scientific Study of Self-Experience with Psychedelics. Journal of Consciousness Studies, 25(11-12), 131–154.

Girn, M., Mills, C., Roseman, L., Carhart-Harris, R. L., & Christoff, K. (2020). Updating the dynamic framework of thought: Creativity and psychedelics. Neuroimage, 116726.

Griffiths, R. R., Johnson, M. W., Carducci, M. A., Umbricht, A., Richards, W. A., Richards, B. D., … Klinedinst, M. A. (2016). Psilocybin produces substantial and sustained decreases in depression and anxiety in patients with life-threatening cancer: A randomized double-blind trial. Journal of Psychopharmacology, 30(12), 1181–1197.

Haak, K. V., & Beckmann, C. F. (2020). Understanding brain organisation in the face of functional heterogeneity and functional multiplicity. Neuroimage, 220, 117061.

Haak, K. V., Marquand, A. F., & Beckmann, C. F. (2018). Connectopic mapping with resting-state fMRI. Neuroimage, 170, 83–94.

Hong, S.-J., de Wael, R. V., Bethlehem, R. A., Lariviere, S., Paquola, C., Valk, S. L., … Smallwood, J. (2019). Atypical functional connectome hierarchy in autism. Nature communications, 10(1), 1022.

Huntenburg, J. M., Bazin, P.-L., Goulas, A., Tardif, C. L., Villringer, A., & Margulies, D. S. (2017). A systematic relationship between functional connectivity and intracortical myelin in the human cerebral cortex. Cerebral Cortex, 27(2), 981–997.

Huntenburg, J. M., Bazin, P.-L., & Margulies, D. S. (2018). Large-scale gradients in human cortical organization. Trends in Cognitive Sciences, 22(1), 21–31.

Johnson, M. W., Garcia-Romeu, A., Cosimano, M. P., & Griffiths, R. R. (2014). Pilot study of the 5-HT2AR agonist psilocybin in the treatment of tobacco addiction. Journal of Psychopharmacology, 28(11), 983–992.

Johnson, M. W., Hendricks, P. S., Barrett, F. S., & Griffiths, R. R. (2019). Classic psychedelics: An integrative review of epidemiology, therapeutics, mystical experience, and brain network function. Pharmacology & therapeutics, 197, 83–102.

Johnson, M. W., Richards, W. A., & Griffiths, R. R. (2008). Human hallucinogen research: guidelines for safety. Journal of Psychopharmacology.

Kraehenmann, R., Pokorny, D., Vollenweider, L., Preller, K. H., Pokorny, T., Seifritz, E., & Vollenweider, F. X. (2017). Dreamlike effects of LSD on waking imagery in humans depend on serotonin 2A receptor activation. Psychopharmacology, 234(13), 2031–2046.

Larivière, S., Vos de Wael, R., Hong, S.-J., Paquola, C., Tavakol, S., Lowe, A. J., … Bernhardt, B. C. (2019). Multiscale Structure–Function Gradients in the Neonatal Connectome. Cerebral Cortex.

Lebedev, A. V., Kaelen, M., Lövdén, M., Nilsson, J., Feilding, A., Nutt, D., & Carhart□Harris, R. (2016).LSD□induced entropic brain activity predicts subsequent personality change. Human Brain Mapping.

Lord, L.-D., Expert, P., Atasoy, S., Roseman, L., Rapuano, K., Lambiotte, R., … Kringelbach, M. L. (2019). Dynamical exploration of the repertoire of brain networks at rest is modulated by psilocybin. Neuroimage, 199, 127–142.

Luppi, A. I., Carhart-Harris, R. L., Roseman, L., Pappas, I., Menon, D. K., & Stamatakis, E. A. (2021). LSD alters dynamic integration and segregation in the human brain. Neuroimage, 227, 117653.

Margulies, D. S., Ghosh, S. S., Goulas, A., Falkiewicz, M., Huntenburg, J. M., Langs, G., … Petrides, M. (2016). Situating the default-mode network along a principal gradient of macroscale cortical organization. Proceedings of the National Academy of Sciences, 113(44), 12574–12579.

Mckeown, B., Strawson, W. H., Wang, H.-T., Karapanagiotidis, T., de Wael, R. V., Benkarim, O., …McCall, C. (2020). The relationship between individual variation in macroscale functional gradients and distinct aspects of ongoing thought. Neuroimage, 220, 117072.

Mesulam, M. (1998). From sensation to cognition. Brain: A journal of neurology, 121(6), 1013–1052.

Millière, R. (2017). Looking for the Self: Phenomenology, Neurophysiology and Philosophical Significance of Drug-induced Ego Dissolution. xFrontiers in Human Neuroscience, 11, 245.

Müller, F., Dolder, P. C., Schmidt, A., Liechti, M. E., & Borgwardt, S. (2018). Altered network hub connectivity after acute LSD administration. NeuroImage: Clinical, 18, 694–701.

Murphy, C., Jefferies, E., Rueschemeyer, S.-A., Sormaz, M., Wang, H.-t., Margulies, D. S., & Smallwood, J. (2018). Distant from input: Evidence of regions within the default mode network supporting perceptually-decoupled and conceptually-guided cognition. Neuroimage, 171, 393–401.

Nichols, D. E. (2016). Psychedelics. Pharmacological Reviews, 68(2), 264–355.

Niendam, T. A., Laird, A. R., Ray, K. L., Dean, Y. M., Glahn, D. C., & Carter, C. S. (2012). Meta-analytic evidence for a superordinate cognitive control network subserving diverse executive functions. Cognitive, Affective, & Behavioral Neuroscience, 12(2), 241–268.

Nour, M. M., & Carhart-Harris, R. L. (2017). Psychedelics and the science of self-experience. The British Journal of Psychiatry, 210(3), 177–179.

Nour M. M., Evans, L., Nutt, D., & Carhart-Harris R. L. (2016). Ego-dissolution and psychedelics: Validation of the ego-dissolution inventory (EDI). Frontiers in Human Neuroscience, 10, 269.

Nutt, D. J., King, L. A., & Nichols, D. E. (2013). Effects of Schedule I drug laws on neuroscience research and treatment innovation. Nature Reviews Neuroscience, 14(8), 577–585.

Paquola, C., Bethlehem, R. A., Seidlitz, J., Wagstyl, K., Romero-Garcia, R., Whitaker, K. J., …Margulies, D. S. (2019). Shifts in myeloarchitecture characterise adolescent development of cortical gradients. Elife, 8.

Paquola, C., De Wael, R. V., Wagstyl, K., Bethlehem, R. A., Hong, S.-J., Seidlitz, J., … Margulies, D. S. (2019). Microstructural and functional gradients are increasingly dissociated in transmodal cortices. PLoS biology, 17(5), e3000284.

Paquola, C., Seidlitz, J., Benkarim, O., Royer, J., Klimes, P., Bethlehem R. A., … Frauscher, B. (2020). The cortical wiring scheme of hierarchical information processing. bioRxiv.

Park, B.-y., de Wael, R. V., Paquola, C., Larivière, S., Benkarim, O., Royer, J., … Valk, S. L. (2020). Signal diffusion along connectome gradients and inter-hub routing differentially contribute to dynamic human brain function. Neuroimage, 224, 117429.

Pokorny, T., Duerler, P., Seifritz, E., Vollenweider, F. X., & Preller, K. H. (2020). LSD acutely impairs working memory, executive functions, and cognitive flexibility, but not risk-based decision-making. Psychological medicine, 50(13), 2255–2264.

Preller, K. H., Burt, J. B., Ji, J. L., Schleifer, C. H., Adkinson, B. D., Stämpfli, P., … Murray, J. D. (2018). Changes in global and thalamic brain connectivity in LSD-induced altered states of consciousness are attributable to the 5-HT2A receptor. Elife, 7, e35082.

Preller, K. H., Duerler, P., Burt, J. B., Ji, J. L., Adkinson, B., Stämpfli, P., … Murray, J. D. (2020). Psilocybin induces time-dependent changes in global functional connectivity: Psi-induced changes in brain connectivity. Biological psychiatry.

Preller, K. H., Herdener, M., Pokorny, T., Planzer, A., Kraehenmann, R., Stämpfli, P., … Vollenweider F. X. (2017). The fabric of meaning and subjective effects in LSD-induced states depend on serotonin 2A receptor activation. Current Biology, 27(3), 451–457.

Preller, K. H., & Vollenweider, F. X. (2016). Phenomenology, structure, and dynamic of psychedelic states. In Behavioral Neurobiology of Psychedelic Drugs (pp. 221-256): Springer.

Roseman, L., Leech, R., Feilding, A., Nutt, D. J., & Carhart-Harris, R. L. (2014). The effects of psilocybin and MDMA on between-network resting state functional connectivity in healthy volunteers. Frontiers in Human Neuroscience, 8.

Schartner, M. M., Carhart-Harris, R. L., Barrett, A. B., Seth, A. K., & Muthukumaraswamy, S. D. (2017).Increased spontaneous MEG signal diversity for psychoactive doses of ketamine, LSD and psilocybin. Scientific Reports, 7, 46421.

Schmid, Y., Enzler, F., Gasser, P., Grouzmann, E., Preller, K. H., Vollenweider, F. X., … Liechti, M. E. (2015). Acute effects of lysergic acid diethylamide in healthy subjects. Biological psychiatry, 78(8), 544–553.

Seeley, W. W., Menon, V., Schatzberg, A. F., Keller, J., Glover, G. H., Kenna, H., … Greicius, M. D. (2007). Dissociable intrinsic connectivity networks for salience processing and executive control. The Journal of Neuroscience, 27(9), 2349–2356.

Sepulcre, J., Sabuncu, M. R., Yeo, T. B., Liu, H., & Johnson, K. A. (2012). Stepwise connectivity of the modal cortex reveals the multimodal organization of the human brain. Journal of Neuroscience, 32(31), 10649–10661.

Setton, R., Mwilambwe-Tshilobo, L., Girn, M., Lockrow, A. W., Baracchini, G., Lowe, A. J., … Nathan Spreng, R. (2021). Functional architecture of the aging brain. bioRxiv, 2021.2003.2031.437922. doi:10.1101/2021.03.31.437922

Studerus, E., Gamma, A., & Vollenweider, F. X. (2010). Psychometric evaluation of the altered states of consciousness rating scale (OAV). PLoS ONE, 5(8), e12412.

Studerus, E., Kometer, M., Hasler, F., & Vollenweider, F. X. (2011). Acute, subacute and long-term subjective effects of psilocybin in healthy humans: a pooled analysis of experimental studies. Journal of Psychopharmacology, 25(11), 1434–1452.

Tagliazucchi, E., Carhart□Harris, R., Leech, R., Nutt, D., & Chialvo, D. R. (2014). Enhanced repertoire of brain dynamical states during the psychedelic experience. Human Brain Mapping, 35(11), 5442–5456.

Tagliazucchi, E., Roseman, L., Kaelen, M., Orban, C., Muthukumaraswamy, S. D., Murphy, K., …Crossley, N. (2016). Increased global functional connectivity correlates with LSD-Induced ego dissolution. Current Biology, 26(8), 1043–1050.

Varley, T., Carhart-Harris, R., Roseman, L., Menon, D., & Stamatakis, E. (2019). Serotonergic Psychedelics LSD & Psilocybin Increase the Fractal Dimension of Cortical Brain Activity in Spatial and Temporal Domains. bioRxiv, 517847.

Vazquez-Rodriguez, B., Liu, Z.-Q., Hagmann, P., & Misic, B. (2020). Signal propagation via cortical hierarchies. bioRxiv.

Vollenweider, F. X., & Preller, K. H. (2020). Psychedelic drugs: neurobiology and potential for treatment of psychiatric disorders. Nature Reviews Neuroscience, 1–14.

Worsley, K. J., Taylor, J., Carbonell, F., Chung, M., Duerden, E., Bernhardt, B., … Evans, A. (2009x). A Matlab toolbox for the statistical analysis of univariate and multivariate surface and volumetric data using linear mixed effects models and random field theory. Paper presented at the NeuroImage Organisation for Human Brain Mapping 2009 Annual Meeting.

Yeo, B. T. T., Kirienen, F. M., Sepulcre, J., Sabuncu, M. R., Lashkari, D., Hollinshead, M., … Buckner, R. L. (2011). The organization of the human cerebral cortex estimated by intrinsic functional connectivity. J Neurophysiol, 106, 1125–1165. doi:10.1152/jn.00338.2011.-

