## Supplementary Figure 1 for "Serotonergic psychedelic drugs LSD and psilocybin reduce the hierarchical differentiation of unimodal and transmodal cortex": Girn-LSD-PSILO_gradients_SUPP_BIORXIV.docx

**Supplementary Methods**

*Participants*

**LSD.** Twenty healthy participants were recruited via word of mouth and provided written informed consent to participate after study briefing and screening for physical and mental health. The screening for physical health included electrocardiogram (ECG), routine blood tests, and urine test for recent drug use and pregnancy. A psychiatric interview was conducted, and participants provided full disclosure of their drug use history. Key exclusion criteria included: < 21 years of age, personal history of diagnosed psychiatric illness, immediate family history of a psychotic disorder, an absence of previous experience with a classic psychedelic drug (e.g. LSD, mescaline, psilocybin/magic mushrooms or DMT/ayahuasca), any psychedelic drug use within 6 weeks of the first scanning day, pregnancy, problematic alcohol use (i.e. > 40 units consumed per week), or a medically significant condition rendering the volunteer unsuitable for the study.

**Psilocybin.** Fifteen healthy participants were recruited via word of mouth and provided written informed consent to participate after study briefing and screening for physical and mental health. The screening for physical health included electrocardiogram (ECG), routine blood tests, and urine test for recent drug use and pregnancy. A psychiatric interview was conducted, and participants provided full disclosure of their drug use history. Key exclusion criteria included: < 21 years of age, pregnancy, personal or immediate family history of psychiatric disorder, substance dependence, cardiovascular disease, claustrophobia, blood or needle phobia, or a significant adverse response to a hallucinogenic drug. All subjects had previous experience with a ‘classic’ psychedelic drug (e.g., LSD, mescaline, psilocybin/magic mushrooms or DMT/ayahuasca) but not within 6 weeks of the study.

*Neuroimaging Data Acquisition*

**LSD.** Neuroimaging data from an already published dataset (Carhart-Harris et al., 2016) was used for the present analyses. The data acquisition and preprocessing details have been described in detail elsewhere (Carhart-Harris et al., 2016); we outline them in brief here. Subjects participated in two scanning days that were separated by 14 days and which each featured three 7-minute resting-state fMRI scans. On a given scanning day, subjects received either a placebo (10ml saline) or LSD (75 µg) via a bolus intravenous injection. The low-moderate LSD dosage was selected to minimize the potential for intra-scanner anxiety while ensuring drug effects (Johnson, Richards, & Griffiths, 2008). The order of the conditions was balanced across participants; participants were blind to this order, but the researchers and those analyzing the data were not. The scans on each of the days were as follows: (1) resting-state eyes-closed with no music, (2) resting-state eyes-closed with music, (3) resting-state eyes-closed with no music. Scans featuring no music (scans 1 and 3) were used in the present analyses.

Resting-state BOLD fMRI data were acquired using a gradient echo planar imaging sequence, TR/TE = 2000/35ms, FoV = 220mm, 64×64 acquisition matrix, parallel acceleration factor = 2, 90◦ flip angle. Thirty-five oblique axial slices were acquired in an interleaved fashion, each 3.4mm thick with zero slice gap (3.4mm isotropic voxels). Structural T1w images were acquired on a 3T GE HDx system. These were 3D fast spoiled gradient echo scans in an axial orientation, with field of view = 256 × 256 × 192 and matrix = 256 × 256 × 20 192 to yield 1mm isotropic voxel resolution. TR/TE = 7.9/3.0ms; inversion time = 450ms; flip angle = 20°.

**Psilocybin.** Neuroimaging data from an already published dataset (Carhart-Harris et al., 2012) was used for the present analyses. The data acquisition and preprocessing details have been described in detail elsewhere (Carhart-Harris et al., 2012) ; we outline them in brief here. Subjects participated in two scanning days that were separated by 14 days and which each featured one 12-minute resting-state scan. Infusion began at 6 minutes following the start of the scan. The post-infusion half of the scan for each condition was used in the present analyses.

Resting-state BOLD fMRI data were acquired using a gradient echo planar imaging sequence, TR/TE 3000/35 ms, field-of-view = 192 mm, 64 × 64 acquisition matrix, parallel acceleration factor = 2, 90° flip angle. Fifty-three oblique- axial slices were acquired in an interleaved fashion, each 3 mm thick with zero slice gap (3 × 3 × 3-mm voxels). A total of 240 volumes were acquired.

**Neuroimaging Data Pre-Processing.** Of the 20 LSD subjects that underwent scanning, 15 were used in the current analyses. One participant was unable to complete the scanning due to anxiety, and four were discarded from analyses due to excessive head motion as measured by framewise displacement (subjects were rejected based on having >15% volumes with FD >= 0.5). This excessive head motion was found in scans conducted during the LSD condition. Of the 15 psilocybin subjects that underwent scanning, 9 were used in the current analyses. Nine were discarded from analyses due to excessive head motion as measured by framewise displacement (subjects were rejected based on having >15% volumes with FD >= 0.5). This excessive head motion was found in scans conducted during the psilocybin condition.

The following pre-processing steps were performed on the BOLD resting-state fMRI data for both datasets: removal of the first three volumes, de-spiking (3dDespike, AFNI), slice time correction (3dTshift, AFNI), motion correction (3dvolreg, AFNI) by registering each volume to the volume most similar to all others, brain extraction (BET, FSL); rigid body registration to anatomical scans, non-linear registration to a 2mm MNI brain (Symmetric Normalization (SyN), ANTS), scrubbing (FD = 0.4), spatial smoothing (FWHM) of 6mm, band-pass filtering between [0.01 to 0.08] Hz, linear and quadratic de-trending (3dDetrend, AFNI), regression of 6 motion parameters, and regression of 3 anatomical nuisance regressors (ventricles, draining veins, and local white matter). Global signal regression was not performed. Quality control tests confirmed the lack of distance-dependent motion confounds in the denoised data (Carhart-Harris et al., 2016).

Structural T1w images were processed using Freesurfer v5.3 ( <http://surfer.nmr.mgh.harvard.edu/>). Structural processing included bias field correction, registration to stereotaxic space, intensity normalization, skull-stripping, and white matter segmentation. A deformable mesh model was fit on the white matter volume via a triangular surface tessellation. This resulted in >160,000 vertices which differentiate gray matter, white matter, and pial surfaces. Individual subject surfaces were fit to the fsaverage5 spherical surface template, which enables stronger inter-subject correspondence in gyral and sulcal folding patterns.

**Subjective Measures.** Subjects completed a number of intra-scanner visual analogue scale (VAS) ratings at the end of each scan for each dataset, reporting on different facets of the psychedelic experience (Carhart-Harris et al., 2016). In addition, subjects completed the 11-factor altered states of consciousness (ASC) questionnaire (Dittrich, 1998; Studerus, Gamma, & Vollenweider, 2010) at the end of each scan day. In the present study, we conducted brain-behaviour analysis with two self-report measures which relate to core components of the psychedelic experience: ego dissolution (“I experienced a disintegration of my 'self' or 'ego”) and complex imagery (a composite of “I could see images from my memory or imagination with exceeding clarity”, “I saw whole scenes in complete darkness or with closed eyes”, and “My imagination was extremely vivid”). The former measure was an intra-scanner VAS rating, while the latter measures were ASC measures conducted at the end of the scan day.

**Supplementary Figures**

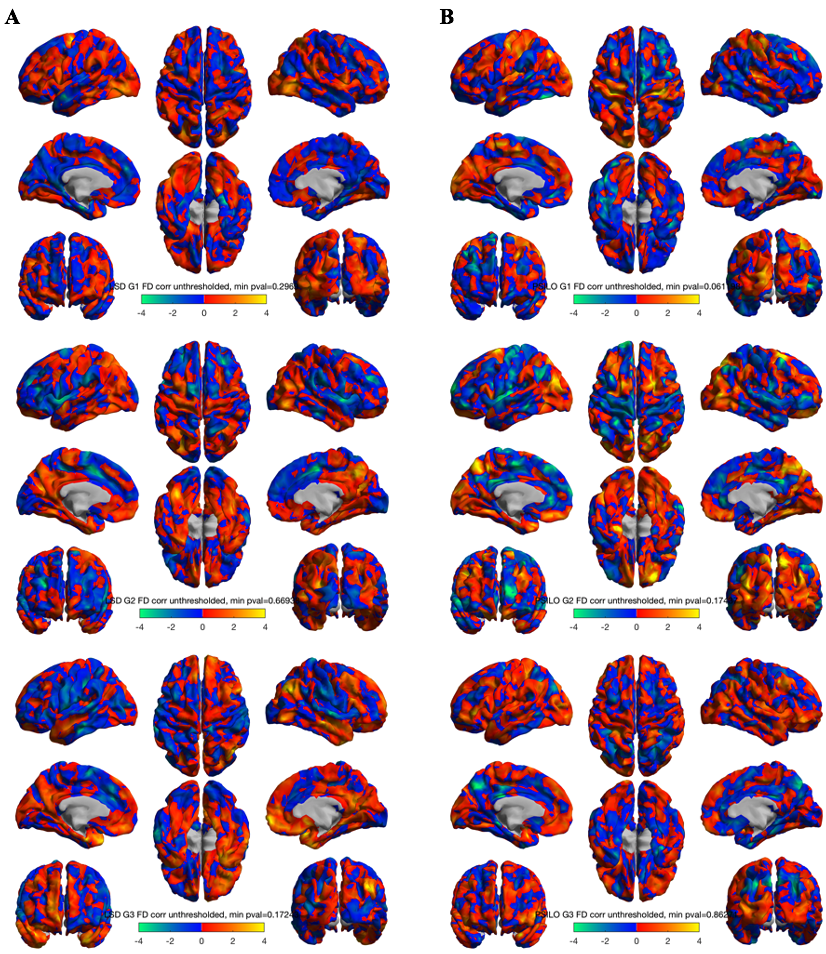

**Supp. Figure 1.** Unthresholded correlations between difference in mean frame-wise displacement and difference in vertex-wise gradient scores, for each of (A) LSD-Placebo and (B) Psilocybin-Placebo contrasts. No significant correlations were found. (Top) principal gradient, (Middle) second gradient, (Bottom) third gradient.

**Supplementary References**

Carhart-Harris, R. L., Muthukumaraswamy, S., Roseman, L., Kaelen, M., Droog, W., Murphy, K., . . . Nutt, D. J. (2016). Neural correlates of the LSD experience revealed by multimodal neuroimaging. *Proceedings of the National Academy of Sciences, 113*(17), 4853-4858. doi:10.1073/pnas.1518377113

Dittrich, A. (1998). The standardized psychometric assessment of altered states of consciousness (ASCs) in humans. *Pharmacopsychiatry*.

Johnson, M. W., Richards, W. A., & Griffiths, R. R. (2008). Human hallucinogen research: guidelines for safety. *Journal of Psychopharmacology*.

Studerus, E., Gamma, A., & Vollenweider, F. X. (2010). Psychometric evaluation of the altered states of consciousness rating scale (OAV). *PLoS ONE, 5*(8), e12412.
